# Leaf abaxial and adaxial surfaces differentially affect plant-fungal pathogen interactions

**DOI:** 10.1101/2024.02.13.579726

**Authors:** Celine Caseys, Anna Jo Muhich, Josue Vega, Maha Ahmed, Aleshia Hopper, David Kelly, Sydney Kim, Matisse Madrone, Taylor Plaziak, Melissa Wang, Daniel J. Kliebenstein

## Abstract

Eudicot plant species have bifacial leaves with each surface varying in a diversity of components, resulting in potentially different microhabitats for pathogens. We tested how *Botrytis cinerea,* a necrotroph fungal pathogen, interacts with the two different leaf surfaces across 16 crop species and 20 Arabidopsis genotypes. This showed that the abaxial surface is generally more susceptible to the pathogen than the adaxial surface. In Arabidopsis, the differential lesion area between leaf surfaces was associated to jasmonic acid (JA) and salicylic acid (SA) signaling and differential induction of defense chemistry. When infecting the adaxial surface, leaves mounted stronger defenses by producing more glucosinolates and camalexin defense compounds, partially explaining the differential susceptibility across surfaces. Testing a collection of 96 *B. cinerea* strains showed genetic heterogeneity of growth patterns, with a few strains preferring the adaxial surface while most are more virulent on the abaxial surface. Overall, we show that leaf-Botrytis interactions are complex with host-specific, surface-specific, and strain-specific behaviors. Within Arabidopsis, this mechanistically links to potential variation in JA/SA signaling across the two surfaces.

## Introduction

When pathogens attack plants, they face a variety of perception/signaling pathways, constitutive and induced physical and chemical defenses that combinatorially control the outcome of the interaction. All these components of plant defense are further influenced by the environmental and genotypic landscapes in which the interaction occurs and vary across plant development. For example, physical and signaling defenses differ between foliar and fruit tissue with fruit ripening altering both forms of defense in plant-pathogen interactions (Forlani *et al*., 2019; Petrasch *et al*., 2019; Silva *et al*., 2023). In leaf, developmental gradients along the axis of the leaf blade linked to response to hormone and growth capacity affect biotrophic fungi such as *Blumeria hordei* (Krasauskas *et al*., 2023). While major developmental transitions have been studied in plant-pathogen interactions, other developmental shifts are less studied. For example, the bifacial surfaces in angiosperm leaves have distinct physical properties yet most infection assays focus on a single leaf surface. The ab- and adaxial leaf surfaces can be considered as different phylloplanes, defined as microhabitats with various micro-topographies hosting various microorganisms. This microhabitat variation raises the question of whether the distinct adaxial (upper) and abaxial (lower) surfaces of the leaf may shift the outcome of the plant-pathogen interaction and how it may vary across both host and pathogen genetic variation.

Studies are beginning to highlight how leaf surface variation affects the host-pathogen outcome. Infections by powdery mildew, an obligate biotroph, are predominant on the adaxial leaf surface (Wu *et al*., 2023). In *Eucalyptus globulus* (blue gum), *Quambalaria eucalyptii* preferentially infected the abaxial leaf surface explained by favorable phylloplane conditions (Mafia *et al*., 2009) while when infected with the necrotroph *Botrytis cinerea*, no surface effect was detected (Caires *et al*., 2014). In hybrids of *Lilium auratum* and *L. speciosum* (oriental lilies), *Botrytis elliptica* failed to invade epidermal cells on the adaxial side while successfully invading from the abaxial (Hsieh *et al*., 2001). Similarly, in *Vicia faba* (faba bean), it was shown that infection from both the specialist *Botrytis fabae* and the generalist *B. cinerea* developed larger lesions on the abaxial surface partly explained by differential induction of a phytoalexin (Hashim *et al*., 1997). While it is becoming clearer that the leaf surface can influence host-pathogen interaction, it remains elusive if this is specifically driven by differences in physical defenses or signaling interactions across the leaf.

The predominant assumption was that variation in the physical properties of the phylloplane leads to altered interactions. The adaxial surface is comprised of the upper epidermis and a dense mesophyll hosting the photosynthetic machinery. On the other side, the abaxial surface is composed of the lower epidermis and loosely packed spongy mesophyll, playing a role in gas exchange and regulation of transpiration. In addition to variation in cell density, the leaf surfaces can also vary in the presence and composition of the cuticle, a thick and rigid chemical barriers made of cutin and waxes (Ziv *et al*., 2018). Trichomes can also provide additional physical barriers preventing fungal spores from reaching the leaf surface or altering water access to the leaf surface. Further, trichomes can anchor growing fungal hyphae close to the leaf surface and contribute to the pathogen propagation (Calo *et al*., 2006). Other surface structures such as stomata and vascular veins influence the micro-topography of the leaf surfaces. The combination of physical properties and architectures suggests that the adaxial and abaxial leaf surfaces may represent different ecosystems with variation in both abiotic conditions such as sunlight and humidity and biotic such as different microbiomes that could influence the interaction (Ritpitakphong *et al*., 2016; Liu *et al*., 2020).

The different leaf surfaces also have different signaling processes and/or outputs that could alter host-pathogen interactions. In leaf bifacial development, auxin signaling plays a major role in controlling complex networks of regulatory genes (Kidner & Timmermans, 2010; Liu *et al*., 2012). The HD-ZIPIII family determines the adaxial while the KANADI and auxin response factor (ARF) families determine the abaxial side. Furthermore, the activity of HD-ZIPIII is negatively regulated by a gradient of microRNAs between the two surfaces. The YABBY family is another important class of transcription factor (TF) involved in both abaxial development and stress responses (Zhang *et al*., 2020). These signaling variations generate expression profiles with hundreds of genes differentially expressed between surfaces (Tian *et al*., 2019). Those differential expression patterns are visible in the defense chemistry with abaxial and adaxial epidermal cells producing varying amounts of metabolites such as flavonols and anthocyanins (Tenorio Berrío *et al*., 2022). In the case of glucosinolates, a family of defense compounds well-described from signaling (Mitreiter & Gigolashvili, 2021) to their impact on plant-pathogen interaction (Plaszkó *et al*., 2022), a tenfold differential concentration was observed in Arabidopsis across the leaf, with more abundance on the abaxial than adaxial surface (Shroff *et al*., 2015).

To begin developing a more general understanding of how leaf surface influences plant-pathogen interactions both across and within plant species, we used *Botrytis cinerea* (grey mold, Botrytis blight, Botrytis thereafter). This generalist fungal pathogen is an endemic pathogen that infects more than a thousand plant species (Fillinger & Elad, 2016; Singh *et al*., 2023). Potentiating Botrytis’ role as one of the top yield loss pathogens (Dean *et al*., 2012), is the polygenic genetic architectures of both Botrytis’ virulence and the hosts’ resistance that influences a wide variety of specific mechanisms (Corwin & Kliebenstein, 2017; Pink *et al*., 2022). Botrytis penetration of the host’s cells can be opportunistic through stomata and wounds or rely on turgor pressure and physical force via the infection cushion to rupture the cuticle and cell wall. Botrytis also secretes cell wall degrading enzymes (Bi *et al*., 2022), toxins (da Silva Ripardo-Filho *et al*., 2023) and small RNAs manipulating the host’s transcriptome (Weiberg *et al*., 2013). Once enough cells are infected and the infection is established, Botrytis’ cell-death inducing proteins contribute to the host’s hypersensitive response, spreading cell death, e.g. the lesion (Veloso & van Kan, 2018; Bi *et al*., 2023). These diverse mechanisms are redundant and build additively based on standing genetic variation (Leisen *et al*., 2022), making Botrytis a fungal pathogen particularly difficult to breed crop defenses against and costly to control. This genetic diversity in mechanisms allows a population of Botrytis isolates to be a highly useful tool in querying how resistance mechanisms vary at different scales.

To assess how Botrytis interactions are influenced by the leaf surface at different evolutionary scales, we tested the effect of leaf surface, across different eudicot species, across different pathogen genotypes and finally across defense mechanisms in a single host species using Arabidopsis mutants. Infecting 16 eudicot species covering 8 families allowed an assessment of potential phylogenetic leaf morphology signals. Using a collection of *A. thaliana* mutants deficient in SA and JA defense signaling, aliphatic and indolic glucosinolates, a putative cyanogenic glycoside, and camalexin biosynthesis, we tested how signaling and chemical defense variation influences the leaf surface effect on the host-pathogen interaction. Finally, we tested the diversity of behaviors of 96 Botrytis strains on the leaf surfaces of the wild type (WT) *A. thaliana* Col-0. This showed that the leaf surface effect on Botrytis-host interactions is genetically conditional across all levels and likely involves both upstream signaling variation and downstream chemical variation but with minimal influence of stomatal content. These experiments highlight the potential necessity to test both leaf surfaces to get more complete plant-pathogen interaction genetic models.

## Material and Method

### Plant growth

Plants for sixteen eudicot species were grown from seeds (Table S1) in a common growth chamber at 20°C with 16 hours daylight. Furthermore, twenty *Arabidopsis thaliana* genotypes and mutants (Miao *et al*., 1994; Cao *et al*., 1997; Zhou *et al*., 1998; Ellis & Turner, 2002; Mikkelsen *et al*., 2003; Barth & Jander, 2006; Sønderby *et al*., 2007; Bu *et al*., 2008; Sonderby *et al*., 2010; Frerigmann & Gigolashvili, 2014; Rajniak *et al*., 2015) were used in this study (Table S2). All Arabidopsis seeds were cold stratified in deionized water for three days at 4°C and sown on soil. After sowing, Arabidopsis plants grew at 20°C with 10 hours photoperiod at 100-120 µE light intensity. All plants were grown for eight weeks in cell-flats containing Sunshine Mix#1 (Sun Gro Horticulture, Agawam, MA) horticulture soil. All plants were watered twice weekly with deionized water for the first two weeks and then with nutrient solution (0.5% N-P-K fertilizer in a 2-1-2 ratio; Grow More 4-18-38).

### Botrytis growth and collection

The isolates for this study were subsampled from a previously described broader collection of 96 *Botrytis cinerea* strains, mostly collected on grapevine in California with about 1/3^rd^ collected around the world on diverse plants (Caseys *et al*., 2021). The strains are maintained by spore collections for long-term preservation as conidial suspension in 60% glycerol at -80^°^C. For each experiment, the strains were grown from spores on peach slices at 21^°^C for two weeks.

### Detached leaf assay

Virulence of the isolates on each species were measured using detached leaf assays (Denby *et al*., 2004). The experiments included four to six replicates (see each experiment section) for each Botrytis strain x leaf surface x host genotype in a randomized complete block design. In brief, mature leaves were cut from the eight weeks old plants and deposited on 1cm of 1% phytoagar either on the abaxial or adaxial surface to provide water to maintain the leaf for the duration of the experiment. Botrytis spores were collected in sterile water, counted with a hemacytometer and sequentially diluted with 50% grape juice to 10 spores/µl.

Drops of 4µl (40 spores of Botrytis) were used to inoculate the leaves. The inoculated leaves were maintained under humidity domes under constant light at room temperature. The experimental trays were pictured every 24 hours. The lesion area on each leaf was measured from the images with a custom-R script (Fordyce *et al*., 2018). In short, images were transformed into hue/saturation/value (hsv) color space. Masks marking the leaves and lesions were created by the script using color thresholds and further confirmed manually. The lesions were measured by counting the number of pixels of the original pictures within the area covered by the lesion mask. The numbers of pixels were converted into centimeter using a reference scale within each image.

### Statistics

All data handling and statistical analyses were conducted in R. The linear modeling was performed with the packages with the lme4 package. The least-square means and standard errors were calculated with the emmeans package using the Satterwaite approximation. Specific models are described within each experimental design section.

### Eudicot experiment

Using the detached leaf assay described above, ten Botrytis strains (Apple517, B05.10, Davis Navel, 2.04.12, Kern B1, Katie Tomato, Pepper, Rose, Triple 3, UKrazz, Table S3) were inoculated on either leaf surface of each host species (Table S1) in a 6-fold replication resulting in 1832 observations. The 10 Botrytis strains were selected to represent a diversity of virulence and host specificity behaviors across eight eudicots species (Caseys *et al*., 2021). The observations were cleaned for failed lesions (technical failures without Botrytis growth) following Caseys et al. 2021, removing 127 observations. Given the different resistance profile of the 16 eudicot species, the lesion areas were centered by subtraction of the mean lesion area of each species. The same method was applied to the stomata density. The least-square mean and standard error were modeled within each species with a linear mixed model: Centered_Lesion∼ Surface + 1|Strain + 1|Tray, where the effect of the Botrytis strains and experimental tray were considered as random effects.

To observe the leaf topology and count stomata, the ad- and abaxial leaf surfaces of the 16 plant species were molded by applying gel cyanoacrylate super glue (the original super glue corporation) on microscope slides and then embedding the leaf on the glue (Castilloa & Ferrarotto, 1998). After drying, the leaf profiles were cleaned in 0.1% anionic detergent solution (Alconox). A Leica CME microscope was used to observe and picture slides.

### Arabidopsis experiment

Using the detached leaf assay described above, ten *B. cinerea* strains (1.03.04, 2.04.17, Apple517, KernB1, Katie Tomato, Pepper, Rasp, Rose, Triple 3, UKrazz, Table S3) were inoculated on either leaf surface of each host species (Table S2) in a 4-fold replication resulting in 1600 observations, from which 107 were considered as technical failures and removed. The 10 Botrytis strains were selected to represent a diversity of virulence and sensitivity to camalexin (Caseys *et al*., 2021). The lesion areas were centered by subtraction of the least-square mean of lesion area of Col-0, the background ecotype for all mutant lines tested. In addition, the Ler ecotype was used to control for whole genome-wide variation. This centering allows directly observing the effect of the gene overexpression or knockout by removing the effect of the genetic background. The surface differential was calculated by subtraction of the least-square mean of lesion area of the adaxial surface to of the least-square mean of lesion area of the abaxial surface. Therefore, a negative surface differential indicates preferential growth on the adaxial surface while positive surface differential indicates preferential growth on the abaxial surface. The least-square mean and standard error were modeled within each genotype with a linear mixed model: Centered_Lesion∼ Surface + 1|Strain + 1|Tray, where the effect of the Botrytis strains and experimental tray were considered as random effects.

To detect the effect of Botrytis on *A. thaliana* aliphatic and indolic glucosinolates, Col-0, *myb28/29* (Sonderby *et al*., 2010), the overexpression *35S:MYB28* (Sønderby *et al*., 2007), *tgg1/2* (Barth & Jander, 2006), *myb34/51* (Frerigmann & Gigolashvili, 2014), *pad3* (Zhou *et al*., 1998), and *cyp79b2/b3* (Mikkelsen *et al*., 2003) genotypes were inoculated on either abaxial or adaxial surface with 40 spores. For each genotype and each surface, infected and uninfected leaves at 72 hours post inoculation (HPI) were transferred into tubes containing 400µl of 90% methanol. The leaves were grinded in the methanol for three minutes using metal beads and a tissue disruptor. After centrifugation, 150µl of the supernatant was transferred on filter plates containing Sephadex DEAE A-25. The first flow through from centrifugation was analyzed for camalexin. Glucosinolates were further washed with 90% methanol and water, before sulfatase incubation overnight (Kliebenstein *et al*., 2001). An injection of 50 µl of Desulfo-glucosinolate solution was run on an Agilent Lichrocart 250–4 RP18e 5 µm column using an Agilent 1100 series HPLC system. Separation was achieved using the following solvent gradient with 1.5 to 5% acetonitrile in 6min, 7% at 8min, 25% at 15min, 92% at 17min and reconditioning of the column in 25min. Glucosinolates were detected with a diode array detector at 229nm. For camalexin, 20µl was injected and separation was achieved with a gradient of aqueous acetonitrile from 31% to 69% acetonitrile in 5min, 50sec to 99% and reconditioning in 8min. Camalexin was detected with a fluorescence detector at emission 318 nm/excitation 385 nm. The area of each infected leaf was measured to normalize the concentrations of glucosinolate (GSL) and camalexin to square centimeter of leaf. Aliphatic glucosinolates are the sum of 3-Methylsulfinylpropyl GSL (3MSO), 4-Methylsulfinylbutyl GSL (4MSO), 4-Methylthio GSL (4MT), 5-Methylsulfinylpentyl GSL (5MSO), 7-Methylsulfinylheptyl GSL (7MSO), and 8-Methylsulfinyloctyl GSL (8MSO). Indolic glucosinolates are the sum of indole-3-yl-methyl GSL(I3M) and 4-methoxy-indole-3-yl-methyl GSL (4MI3M). To estimate the percentage of variance in lesion area, linear modelling across six mutants and Col-0 was performed based chemical content with the formula: Lesion ∼ Isolate*Surface*Camalexin + aliphatic + indolic. Linear regressions between lesion area and camalexin, I3M, 4MI3M or aliphatic glucosinolates were also performed for each genotype in which the targeted compound was detected.

For further condensing the information into a single figure, the data were centered by subtraction of the mean of concentration of the mock (not infected leaves) in each genotype for each compound. This mean-centering allowed us to directly plot the concentration induced by the pathogen on each leaf surface. The data for camalexin was not mean-centered, given that camalexin is not constitutively synthesized in Arabidopsis and uninfected leaves do not produce camalexin. For each compound, a linear mixed model was used to model the least-square means and standard error: concentration∼ Surface + 1|Strain + 1|Tray, where the effect of the Botrytis strains and experimental tray were considered as a random effects.

To detect the mode of infection of Botrytis on *A. thaliana* leaf surfaces, Col-0, *myb28/29* (Sonderby *et al*., 2010), the overexpression *35S:MYB28* (Sønderby *et al*., 2007), *tgg1/2* (Barth & Jander, 2006), *myb34/51* (Frerigmann & Gigolashvili, 2014), *pad3* (Zhou *et al*., 1998), and *cyp79b2/b3 (Mikkelsen et al., 2003)* genotypes were sprayed on either abaxial or adaxial surface with water containing 10 spores/µl. After 24, 48, and 72 hours post inoculation, leaves were cleared with acetic acid – ethanol 1:3 v/v solution for 3 hours. They were then further cleared with acetic acid-ethanol-glycerol 1:5:1 v/v/v solution for another 3 hours. The leaves were then stained with lactic acid-glycerol-water 1:1:1 v/v/v containing 0.05% trypan blue. Leaves were mounted on microscope slides after rinsing in 60% glycerol.

### Botrytis diversity experiment

Using the detached leaf assay described above, we infected Col-0 detached leaves with a 96 strains from Botrytis (Caseys *et al*., 2021) in a 4-fold replication resulting in 792 observations. The 96 strains include the various strains tested in the Eudicot and Arabidopsis experiments mentioned above (Table S3). The lesion area was modeled with a linear model Lesion_Area ∼ Strain + Surface + Strain*Surface + Experiment_Tray + Individual_Plant. Strain and surface are biological factors while experimental tray and the individual plant from which the leaves were collected are technical factors.

## Results

### What is the range of leaf surface effects within Botrytis-eudicot interactions?

To test the evolutionary diversity of potential leaf surface effects on host-Botrytis interactions, we infected 16 eudicot species with 10 diverse strains of Botrytis, chosen to represent the range of virulence and host specificity of the pathogen (Table S3). These plant species were chosen to cover eight families, from the core eudicots, rosids, and asterids (Figure 1). Further, these species sample a diversity of leaf thickness, stomata sizes and densities, and micro-topographies across the leaf surfaces (Table S1, Figure S1). For example, measuring stomatal density across the species confirmed the expected relationship where the abaxial surface has more stomata (range 215-1052 stomata/mm^2^, Table S1) than adaxial surface (range: 0-556 stomata/mm^2^, Table S1). However, this ratio is highly variable with some species like *A. thaliana*, and *Ocimum basilicum* (basil) having a small difference between the surfaces while some like *Phaseolus vulgaris* (bean), *Capsicum annuum* (pepper) and *Mentha spicata* (mint) have near presence/absence variation across the surfaces (Figure 1B). The adaxial surface tends to be flatter than abaxial due to the veins on the abaxial surface that create ridges and valleys. However, the variation of micro-topographies across plant species is further driven by the presence and type of trichome (Figure S1). This confirms that there is a range of physical properties across the species and leaf surfaces.

**Figure 1.**
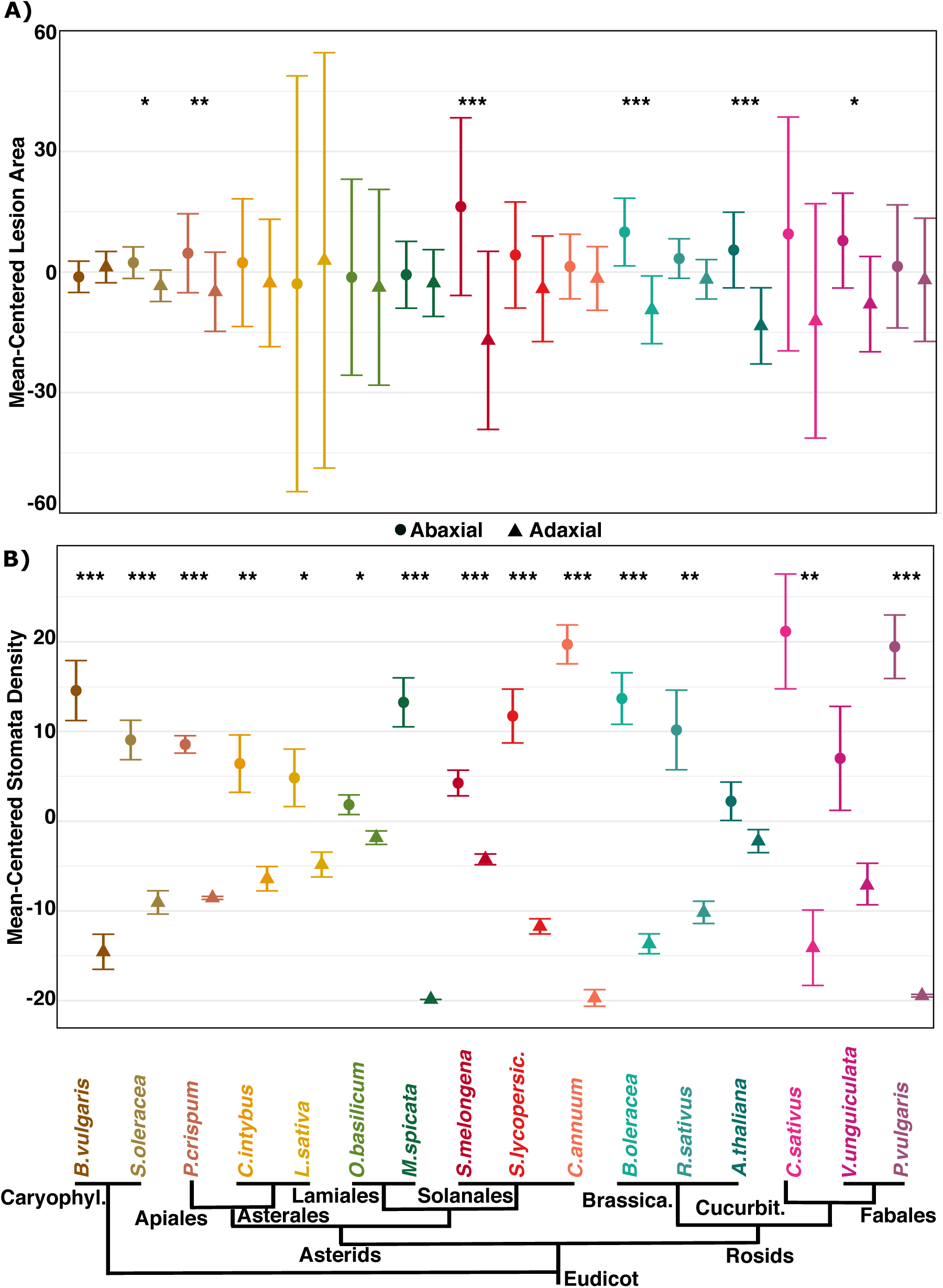
The leaf surface effect of 16 eudicot species on lesion area and stomatal density. Given the large variation among the species in term of resistance and developmental patterns, the data was centered by subtraction of mean of each species to parse the surface effect from the species effect. Circles represent the abaxial surface while up-pointing triangles represent the adaxial surface. Level of significance of within species surface effect: *** p<0.001, ** p<0.01, * p<0.05. A) Mean-centered lesion area [mm2] and standard error bars measured at 72 HPI on each surface. B) Mean-centered stomatal density (stomata/mm^2^; Table S1) and standard error bars on each surface.

Infecting the collection of Botrytis strains on the leaves of all species showed an overall trend for larger lesion areas on the abaxial than adaxial surfaces. Representative raw images of leaf surfaces and lesions are provided in Figure S2. The larger abaxial lesions were measured on all species except *Beta vulgaris* (chard) and *Lactuca sativa* (lettuce) (Table S1, Figure 1A). This trend was statistically significant in six species (Figure 1A) spread across the Eudicot phylogeny; *A. thaliana*, *Vigna unguiculata* (cowpea), *Solanum melongena* (eggplant), *Brassica oleracea* (kale), *Petroselinum crispum* (parsley), and S*pinacia oleracea* (spinach). The variance in standard error was explained by both the host susceptibility (e.g. large lesion have larger standard deviation) and different sensitivity to the microenvironment, such as small variation airflow or light in the experimental tray (Figure S3). Using the physical properties previously measured, we tested how variation in physical properties links to variation in the Botrytis-host interactions across the eudicots. However, the lesion area differential between leaf surfaces had no correlation to stomatal density nor leaf thickness (Figure S4). This suggests that the leaf surface effect on host-Botrytis interactions is variable across the eudicots and is not predominantly driven by these measured physical aspects like stomatal density or leaf thickness variation.

Given the extensive availability of mutants in *A. thaliana*, we utilized mutants that alter defense chemistry and/or defense signaling to test of how they affect Botrytis lesion formation when infected on different leaf surfaces. We collected 18 mutants all within the Col-0 background (Table S2) with Ler as an additional natural accession. These mutants modify the jasmonic acid and salicylic acid signaling, camalexin, aliphatic glucosinolate, and indolic glucosinolate related metabolism, and were infected with the 10 Botrytis strains selected to represent a range of virulence and camalexin sensitivity (Table S3). The level of significance of the surface effect in Col-0 (Figure 1, 2) depended on the set of ten Botrytis strains tested (Table S3).

Measuring the lesion area on the leaf surfaces on these Arabidopsis genotypes with various susceptibilities revealed three patterns. First, some mutants had limited or inversed surface differentials compared to WT, e.g. the size of the lesions on both surfaces were more equal than in WT. This pattern was observed for *cyp81D8*, the double knockout mutant of *myb28/29*, the single knockout of myb29, the *35S:MYB28* overexpression line, and the *35S:MYB29* overexpression line (although *35S:*MYB29 had a strong differential at 72h and then plateaued, Figure 2B). Interestingly, the MYB mutants are known to alter the spatial pattern of aliphatic glucosinolates across the leaf (Malitsky *et al*., 2008; Sonderby *et al*., 2010). In the second observed pattern, the mutations amplified the surface effect indicated as having a significantly larger lesion area on abaxial than adaxial surface without increasing the overall plant susceptibility (Figure 2A). This pattern was observed in *npr1* and *pad4* involved in salicylic acid signaling defense responses. It was also observed *in tgg1/2*, the double mutant for the two myrosinase enzymes that convert glucosinolates into toxic isothiocyanates. The mutants in *cyp71a12* and *cyp82c2* involved in the biosynthesis of 4-hydroxy indole3-carbonyl nitrile, a potential indole-cyanogenic glycoside phytoalexin, also showed this pattern. In the final pattern, the mutants lead to an increased surface effect as well as an increase in overall plant susceptibility (larger lesion area compared to Col-0, Figure 2A). This pattern was observed for *coi1-16* involved in jasmonic acid signaling and *tga3* a TF involved in salicylic acid signaling defense response. The double mutants *myb34/51*, *cyp79b2/cyp79b3* and the *pad3* single mutant, all abolishing camalexin production also showed this pattern. While the surface differential in lesion area largely increases in a linear fashion in time as with Col-0, some mutant lines have different temporal trajectories including the biosynthetic mutants *cyp79b2b3*, *cyp81D8* and 35S:*MYB29* (Figure 2B). This suggests that the differential surface effect is affected by both signaling and defense metabolism pathways in Arabidopsis-Botrytis interactions.

**Figure 2.**
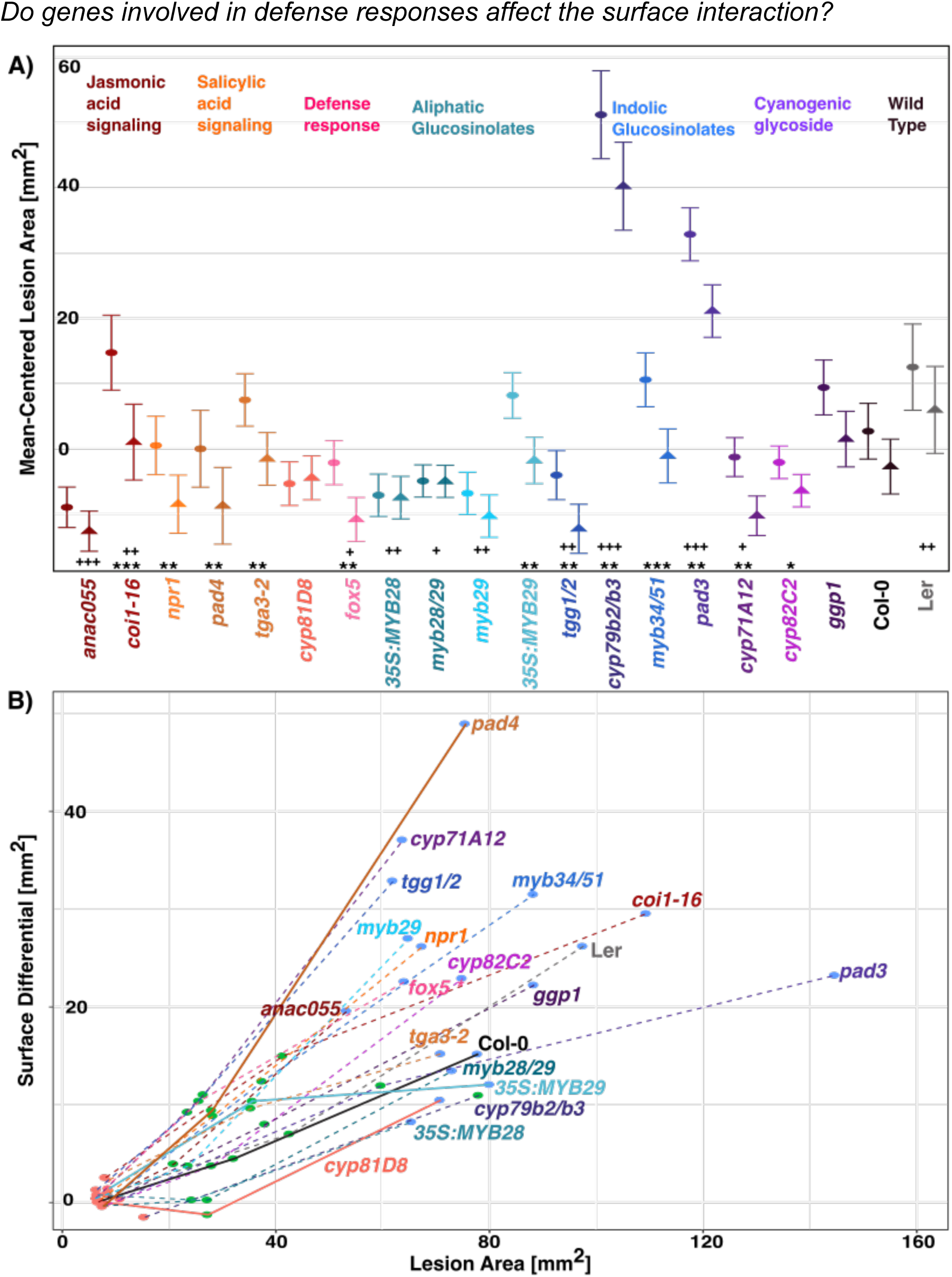
The leaf surface effect of *A. thaliana* mutant lines colored by pathways, the Col-0 background and a wild-type control, Ler. A) Lesion area [mm^2^] centered on Col-0 (background) and standard error bars as modeled at 72 HPI on each surface. To parse the effects of the background ecotype, lesion areas were centered by subtraction of the least-square mean of lesion area of Col-0. The abaxial surface is represented by circles while the adaxial surface by up-pointing triangles. Level of significance of surface effects within each genotype: *** p<0.001, ** p<0.01, * p<0.05. Level of significance of pairwise ANOVA that compare each genotype to Col-0: **^+++^** p<0.001, **^++^** p<0.01, **^+^** p<0.05. B) Lesion area and surface differential (abaxial-adaxial lesion area) at 48 HPI (red dots), 72 HPI (green dots) and 96 HPI (blue dots). The high susceptibility double mutant *cyp79b2/b3* is only traced up to 72 HPI, because at 96 HPI the entire Arabidopsis leaf area is consumed.

To test if Botrytis’ early growth and infection behavior/strategy differs between the two leaf surfaces and if this is altered by the mutants, we infected, stained and microscopically observed Botrytis hyphae on leaf surfaces from Col-0, *myb28/29, 35S:MYB28, tgg1/2, myb34/51, pad3, and cyp79b2/b3,* mutants chosen for their range of effect (Figure 2). In this analysis, we did not observe Botrytis penetration through stomata, but instead the infection cushions, a multicellular structure specialized in breaching the host tissue through mechanical and chemical actions, appeared to directly target pavement cells (Figure S5). The same behavior was seen across all samples including abaxial and adaxial leaf surfaces and all genotypes. These observations reinforce the lack of correlation between lesion area and stomatal density observed in the eudicots (Figure S4) while suggesting that the differential lesion area observed between leaf surfaces is not associated with a dramatic shift in Botrytis tissue penetration strategy.

### Metabolic responses to estimate surface signaling differences

Ten Botrytis strains were infected on the abaxial and adaxial surface of six defense metabolite mutants (*35S:MYB28, myb28/29, tgg1/2, myb34/51, cyp79b2/b3* and *pad3*) and their wild-type genotype Col-0. To estimate the role of the parameters tested (e.g. Botrytis stains, leaf surfaces, variation in chemical defenses across mutants) in the lesion area, we performed linear modeling. This model revealed that the diverse behavior of the ten Botrytis strains explained the largest proportion (39.1%, Figure S6) of the variance in lesion area. The chemical defenses induced by the infection also had a large effect. The concentration of camalexin explained 15.7% while the concentrations of indolic and aliphatic GSLs explained respectively 3.2% and 1.1% of the variance in lesion area (Figure S6). The effect of infecting the abaxial and adaxial leaf surfaces was smaller, explaining 2.5% of the variance in lesion area, but also contributed in interaction with other parameters such as camalexin (Figure S6). To further understand the effect of each chemical in the defense metabolism mutants, we further analyzed by linear regression their relation to lesion area. This analysis revealed negative association to lesion area of camalexin in *35S:MYB28, myb28/29, tgg1/2,* Col-0 (Figure S7), I3M in *tgg1/2, pad3,* Col-0 (Figure S8) and aliphatic GSLs in *myb34/51, cyp79b2/b3* and Col-0 (Figure S10). These results confirmed and quantified the defensive role of induction of camalexin and glucosinolates by the plant to fight Botrytis.

Given that defense metabolites mutants shifted the leaf surface interaction (Figure 2) and are regulated by both SA and JA pathways, these mutants can be used as a proxy measure to estimate differential signaling across infected leaf surface. Interestingly, in the WT controls, there was no statistical support for differential induction depending on the leaf surface that was infected (Figure 3). The mutants however identified differential induction linked to infection based on leaf surface. For example, mutants in the aliphatic glucosinolate TFs (myb28/29) lead to a change in the camalexin response (Figure S7) with a higher response to adaxial than abaxial infection (Figure 3A). Mutants in the myrosinases (tgg1/2) and the camalexin biosynthetic gene (pad3), led to shifts in indole glucosinolate responses (Figure S8, S9), again with a higher adaxial than abaxial response (Figure 3BC). This shows that there are differential chemical responses to adaxial versus abaxial infection. Interestingly, the mutants altered the response of indirectly related chemicals, e.g. aliphatic mutants altered indolic responses and vice versa. This indicates that both the SA/JA signaling pathways and the metabolites they control likely influence part of the difference in the host-pathogen interaction across the two leaf surfaces. Future work will be needed to uncover the specific regulatory mechanisms and how they vary across leaf surfaces leading to these differences.

**Figure 3:**
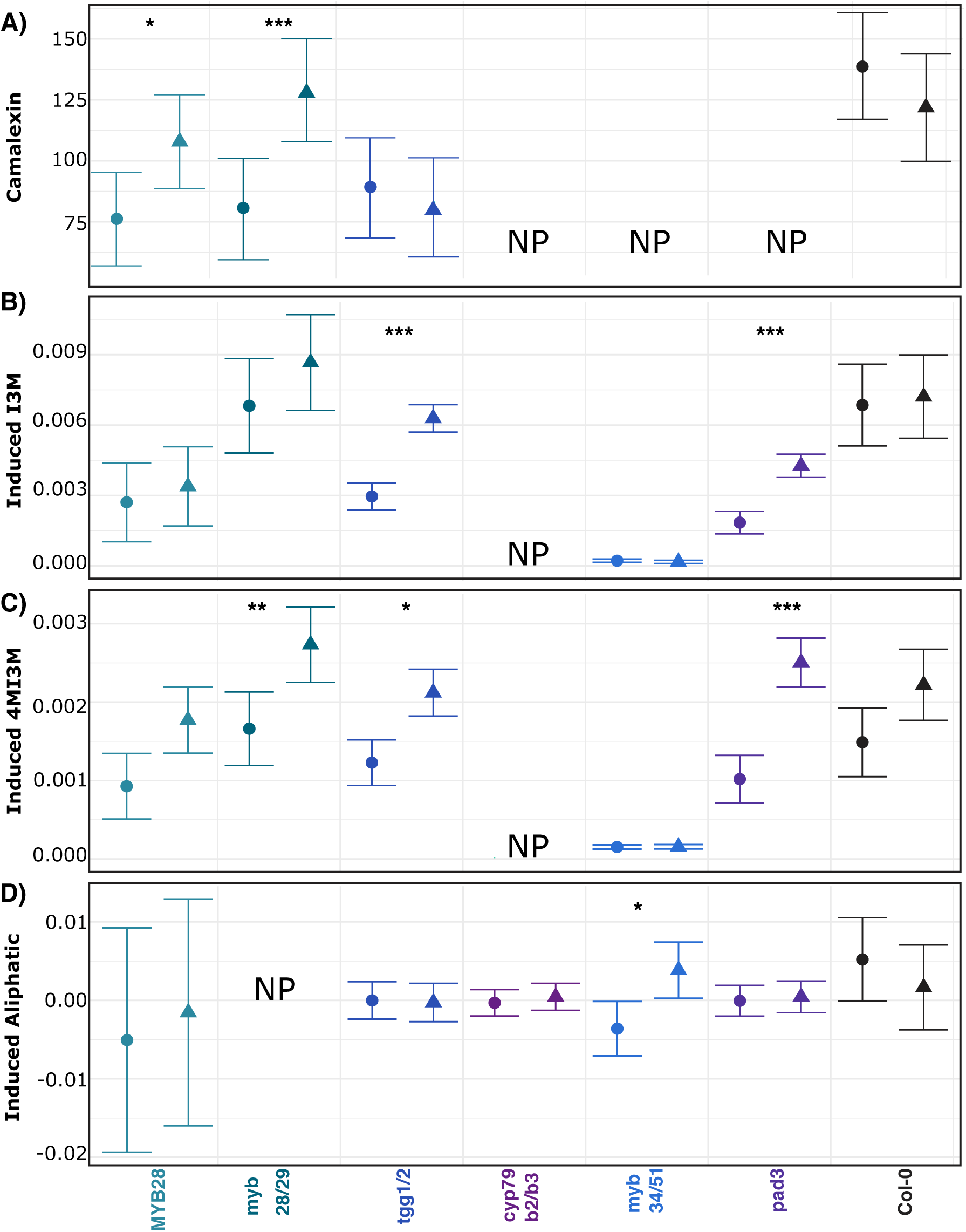
Camalexin, induced indolic, and aliphatic glucosinolates at 72 HPI of either abaxial or adaxial leaf surface with *B. cinerea*. Induced indolic and glucosinolates levels are centered by subtraction of the mean of concentration of the uninfected leaves. The abaxial surface is represented by circles while the adaxial surface by up-pointing triangles. Level of significance: *** p<0.001, ** p<0.01, * p<0.05. A) Mean and standard error bars of the camalexin in nmol/cm^2^. A) Mean and standard error bars of the indol-3-yl-methyl glucosinolate (I3M) in nmol/cm^2^. C) Mean and standard error bars of the 4-methoxy-indole-3-yl-methyl glucosinolate (4MI3M) in nmol/cm^2^. D) Mean and standard error bars of aliphatic glucosinolates in nmol/cm^2^. ABCD) NP stands for “not present”. The mutants *cyp79b2/b3, myb34/51* (Schlaeppi *et al*., 2010) and *pad3* (Zhou *et al*., 1998) do not produce camalexin. The mutant *cyp79b2/b3* doesn’t produce indolic glucosinolates while *myb34/51* does not fully stop their synthesis, probably due to accessory role of MYB122 (Henning & Tamara, 2014). The mutant *myb28/29* doesn’t produce aliphatic glucosinolates (Sonderby *et al*., 2010).

### How does B. cinerea genetic diversity influence growth on leaf surfaces?

The previous experiments show that the genetic variation both between and within eudicot species can influence the effect of leaf surface on Botrytis virulence. Previous work has shown extensive genetic variation within Botrytis with regards to phytochemical detoxification and cell wall degrading proteins that could also influence the leaf surface interaction. This leads us to test how genetic variation in the pathogen can influence the leaf surface effect. To do this, we infected Col-0 with a collection of 96 strains of Botrytis and tracked the lesion development on leaf surfaces.

To measure the contribution of the interaction of the genetic diversity in the host and the pathogen into lesion development, we modeled the growth of the lesion area on the leaf surfaces with a linear model at each time point. The linear models revealed that the diversity of strains accounts for at least 35% of the variance (Figure 4) that is maximal around 72 HPI. The variance due to abaxial and adaxial leaf surfaces increased from 3% at 48 HPI to 20% of the total variance in lesion area at 96 HPI (Figure 4B). As the genetic diversity in the strain collection is a dominant component controlling the leaf-surface interaction, we investigated the 96 strains for their virulence and surface differential (lesion area on the abaxial – adaxial surface) across the three time points. The relationship between virulence and surface differential is not consistently linear as various growth patterns are present in the strain collection (Figure 4C). Most strains develop consistently faster on the abaxial surfaces (e.g. strain 1.05.22, Figure 4D) while some stabilize their growth on both surfaces after 72 HPI (e.g. strain 2.04.12, Figure 4D). Other strains have high virulence but low preference for growth on a surface (e.g. strain 2.04.14, Figure 4D). Finally, few strains indicate a degree of preference to grow on the adaxial surface (e.g. strain 2.04.04, Figure 4D). These various growth patterns suggest that virulence and surface effect require different sets of genes in Botrytis.

**Fig. 4:**
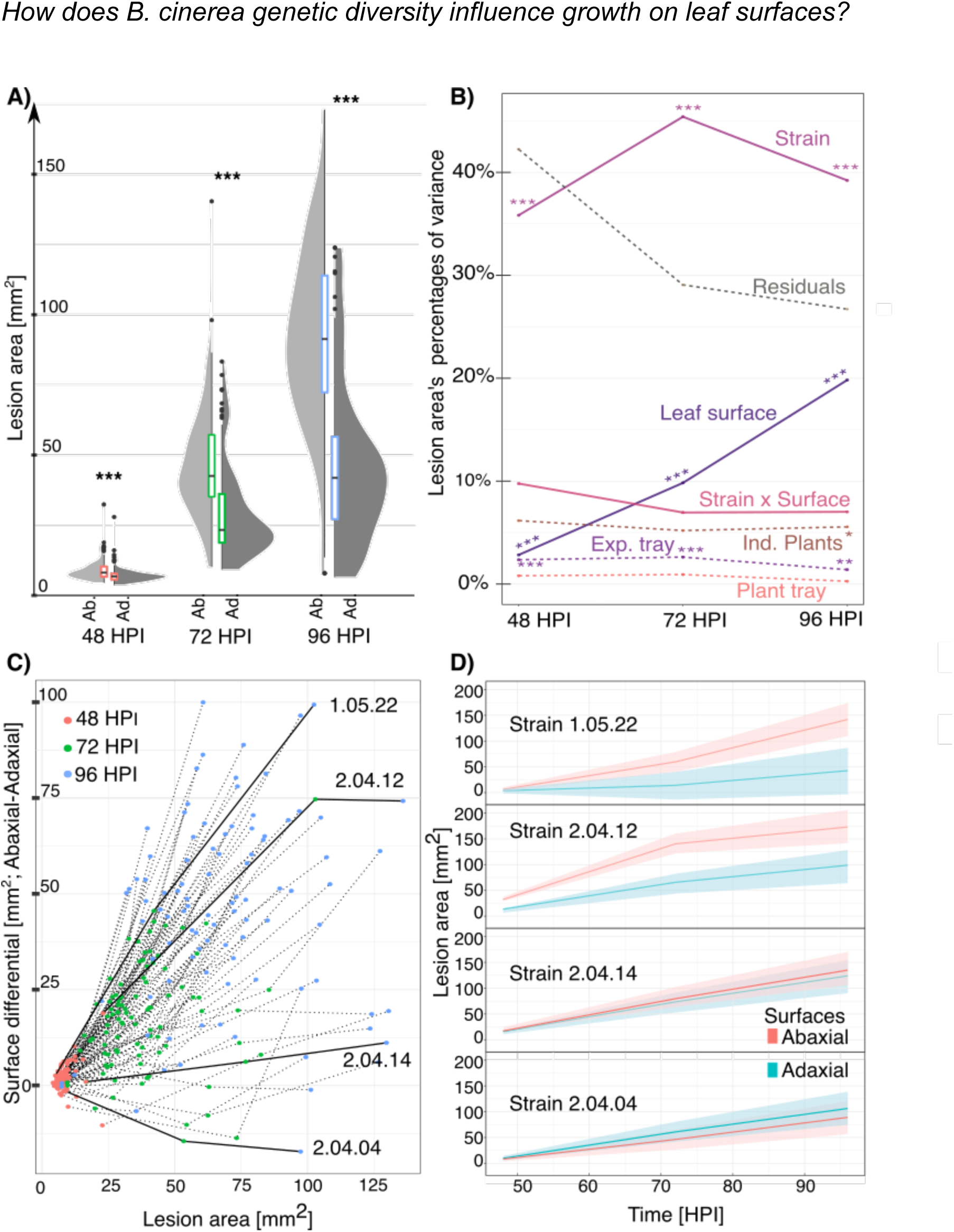
Growth of 96 B. cinerea strains on A. thaliana (Col-0) abaxial and adaxial surfaces. A) Violin and boxplot of lesion area of at 48 HPI (in red), 72 HPI (in green), and 96 HPI (in blue) for 96 Botrytis strains. B) Percentage of variances explained of seven biological and technical parameters calculated by linear modeling of the lesion area on Col-0 leaves at 48, 72 and 96 HPI. Plain lines represent the biological factors while dashed lines represent technical factors and residuals. Significance: *** p<0.001; ** p<0.01; * p<0.05. C) Lesion development of 96 Botrytis strains represented by the mean of lesion area across both surfaces (x-axis) and the difference in lesion area between the abaxial and adaxial leaf surface (y-axis). D) Estimated marginal means and 95% confidence interval of the lesion area of the four strains highlighted in panel C.

## Discussion

By surveying Botrytis virulence across different evolutionary scales, we show that developmental variation across the adaxial and abaxial leaf surfaces influence the interaction with Botrytis across diverse eudicot host species. The difference in lesion area between adaxial and abaxial surfaces differed across the tested eudicot species and was not predictable based on the classically assumed physical properties such as variation in stomatal density and leaf thickness. It remains to be seen if other unmeasured physical or microbiome components could be critical to this variation, especially in interaction with diverse strains of Botrytis (Rowe & Kliebenstein, 2010).

As shown by testing 96 strains, *B. cinerea* as a species encompasses many different growth patterns when interacting with the leaf surfaces (Fig.4). Whether that complex interaction is a sum of independent effects in the host and strain (Plant susceptibility + Leaf Surface + Strain virulence) or whether those effects interact (Plant susceptibility * Leaf Surface * Strain virulence) remain to be determined. Given the wide range of hosts (Singh *et al*., 2023) and organs attacked by Botrytis and the polygenic nature of its virulence, it is challenging to know whether those growth patterns might results from genes’ presence/absence variation (Simon *et al*., 2022), single nucleotide polymorphisms (Caseys *et al*., 2021) or plastically controlled regulation of virulence pathways. Furthermore, how various strains detect the hosts’ extracellular signals and mount diverse responses/strategies at different times post inoculation remains largely unknown.

Botrytis has over a thousand surface-associated proteins that might contribute to detecting the host (Escobar-Niño *et al*., 2021), with signal transduction best described through the interplay of cAMP, MAP-kinase, histidine-kinase and calcium-dependent signaling pathways (Liñeiro *et al*., 2016). Even in absence of precise mechanisms, it remains striking how complex the host-Botrytis interaction is with developmental patterns such as the leaf surface driving large variation in lesion area.

Using Arabidopsis mutants that altered differently the SA and JA defense signaling pathways began to illuminate some of the mechanisms involved. Once the pathogen is detected on a leaf surface and immunity is triggered, defense response signals are integrated into complex networks, with SA and JA signaling in competitive intercommunications. This complex signal integration is beneficial to canalize the range of virulence both within and between pathogens species by fine-tuning the immune response (Caarls *et al*., 2015; Zhang *et al*., 2017; Hou & Tsuda, 2022). Interestingly, the Arabidopsis experiment showed that the SA signaling mutants (*pad4, npr1*) have strong surface effects without increasing the overall plant susceptibility while JA signaling (*coi1, anac055*) strongly impacted the plant susceptibility (Figure 2b)(Rowe *et al*., 2010). This suggests that SA/JA canalization contribute not only to fine-tune the immune response to the pathogens’ virulence but also to influence developmental signals. Such spatiotemporal developmental patterns were also recently described in Barley with variation in response to JA and gibberellins across the leaf blade infected by *Blumeria hordei (Krasauskas et al., 2023).* In Arabidopsis leaves infected with powdery mildew, higher resistance of the abaxial surface is explained by defenses associated to both SA pathway and glucosinolates (Wu *et al*., 2023). In Botrytis, the plasticity of signal integration is not only strain-dependent (Zhang *et al*., 2017; Zhang *et al*., 2019) but also developmentally driven and time-dependent as those patterns increased with time post inoculation (Figure 2b). Proceeding to Arabidopsis mutants that alter the outputs from the SA/JA pathways allowed finer scale assessment of mechanisms influencing differential leaf surface interactions. From our experiment, the role of MYBs controlling defense metabolism altered the surface pattern that emerged with indolic associated MYB34 and MYB51 increasing the difference, aliphatic glucosinolate MYB28 diminishing the surface effect and the other aliphatic MYB having a time dependent effect. This suggests that the defense signaling controlled by these MYBs varies across infection of the leaf surfaces, especially linked to specialized metabolites, as demonstrated by the strong surface differential in cyp79b2/b3, cyp71a12, cyp82c2 and tgg1/2. Those patterns suggest surface-dependent gene expression of both MYBs and secondary metabolite enzymes, differentials that are also suggested by existing datasets. From the shoot apical meristem atlas at early stage of bifacial leaf development (Tian *et al*., 2019), among the genes we tested, CYP81D8 and MYB51 are preferentially expressed on the abaxial while MYB28, MYB29, and MYB34 are preferentially expressed on the adaxial domain. In the Arabidopsis single-cell dataset (Tenorio Berrío *et al*., 2022), adaxial and abaxial pavement cells were identified, revealing abaxial focused expression of GGP1, a key enzyme in both glucosinolate and camalexin synthesis. Additionally, these MYBs are known to have altered expression in Arabidopsis mutants with altered ad-abaxial development supporting their potential role in differentially influencing virulence across leaf surfaces (Malitsky *et al*., 2008).

Further support for altered signaling was obtained by quantifying glucosinolates and camalexin in six mutant lines infected on one or the other surface. This showed differential induction of defense compounds with infection on the adaxial surface (Figure 3). The localization of glucosinolates within leaves under infection and whether this results from de novo synthesis at the site of the attack or strategic movement of defenses within the leaf remains largely unknown (Burow & Halkier, 2017). Though differential induction of indolic glucosinolates and camalexin might contribute to the surface effect, these defense compounds are restricted to the Brassicaceae. Given that the surface effect is spread across the eudicots, other phytoalexins might have similar differential induction in other eudicot species. This is similar to the observation that most lineage specific phytoalexins and specialized metabolites are routinely controlled by the conserved JA/SA signaling pathways (Lacchini & Goossens, 2020; Rieseberg *et al*., 2023).

While the complexity of how abaxial and adaxial surfaces regulate genes and metabolites is only starting to be addressed, how those react to a pathogen infection remains largely unaddressed. Single-cell sequencing of Arabidopsis infected by the biotroph fungal pathogen *Colletotrichum higginsianum* revealed the spatially dynamic and cell-type specific plant response (Tang *et al*., 2023). Although the fungal infections were inoculated only on the adaxial surface, they especially show the crucial role of intracellular immune receptors (NLRs) across cell types and cell-specific induction of indolic glucosinolates. Future experiments will need to confirm the differential expression of immune receptors and other defense components not only across cell types but also along the ab-adaxial axis.

To conclude, we provided a snapshot into how variable plant-pathogen interactions are from large scale (e.g. across the plant kingdom) to small scale (e.g. leaf surfaces). We also highlight a potential ascertainment bias in foliar infection studies: most infection assays to validate genes and determine mechanisms are necessarily done on a single surface, typically adaxial, to make the experiments feasible. However, the pathogens or plant mutants/genotypes may show differential behavior if infected on the abaxial surface. To determine precisely the signaling cascade and gene networks responsible for the larger Botrytis lesions on the abaxial surface will require both the host and pathogen transcriptomes to be analyzed in detail with single-cell experiments in three dimensions across the time of the infection. Conducting such an experiment is currently limited by available technology, such as the absence of a cell atlas for Botrytis. The width of the leaf will need to be measured in its entirety to allow for cell types to be fully differentiated by their location on the abaxial-adaxial axes, revealing for example whether the defense chemicals are produced locally or relocated from storage. Such limitations might be lifted by new technologies that will preserve the special context of tissues (Nolan & Shahan, 2023).

## Acknowledgments

This work was initiated by a curiosity-driven question of an undergraduate student asking why we inoculated only the adaxial surface. Undergraduate students also largely conducted the experiments and observations. This work was supported by the NSF award IOS 2020754 and USDA award 2019-05709 to DJK.

## Authors contribution

Conceptualization: C.C., D.J.K; Supervision: D.J.K.; Data analysis: C.C.; Funding acquisition: D.J.K., Investigation: C.C., A.J.M., J.V., M.A., A.H., D.K., S.K., M.M., T.P.,M.W.; Writing: C.C., A.J.M and D.J.K.

## Data availability

Correspondence and requests for materials should be addressed to.

