## Supplemental Figures and tables for "Leaf abaxial and adaxial surfaces differentially affect plant-fungal pathogen interactions"

| | |
|---|---|
| 0000-0003-4187-9018 | Celine Caseys |
| 0000-0001-5759-3175 | Daniel Kliebenstein |
| 0000-0002-0262-1252 | Melissa Wang |


**Keywords:** plant-pathogen interactions, Botrytis cinerea, necrotroph fungi, crops, leaf surfaces, defense signalling, plant development

### Supplemental Material

| Phylum | Family | Species | Common name | Genotype | Company | Average Leaf Thickness [uM] | Stomata per mm2 Adaxial | Stomata per mm2 Abaxial | Lesion area Adaxial | Lesion area Abaxial |
|---|---|---|---|---|---|---|---|---|---|---|
| Core Eudicot | Caryophyllales | *Spinacia oleracea* | Spinach | Bloomsdale longstanding | Cornucopia seeds | 323 | 236.31 | 491.39 | 22.08 | 27.94 |
| Core Eudicot | Caryophyllales | *Beta vulgaris* | Swiss chard | Fordhood Giant | Cornucopia seeds | 371 | 190.92 | 600.94 | 27.23 | 25.07 |
| Asterids | Lamiales | *Ocimum basilicum* | Basil | Italian Genovese | Botanical interests, Inc | 226 | 170.58 | 222.22 | 116.66 | 122.45 |
| Asterids | Lamiales | *Mentha spicata* | Mint | | Botanical interests, Inc | 260 | 0.00 | 466.35 | 46.91 | 49.29 |
| Asterids | Solanales | *Solanum lycopersicum* | Tomato | Large red cherry | Green garden products | 324 | 61.03 | 391.24 | 87.29 | 98.39 |
| Asterids | Solanales | *Solanum melongena* | Eggplant | Black Beauty | Green garden products | 310 | 96.24 | 215.96 | 99.18 | 127.71 |
| Asterids | Solanales | *Capsicum annuum* | Pepper | Felicity | Territorial seeds | 277 | 318.31 | 873.24 | 26.98 | 32.19 |
| Asterids | Asterales | *Lactuca sativa* | Lettuce | Chinese stem (LJ10335) | Warwick University | 120 | 98.59 | 234.74 | 77.54 | 83.08 |
| Asterids | Asterales | *Cichorium intybus* | Chicory | Augusto (PI.651886) | USDA GRIN | 232 | 136.15 | 316.90 | 75.19 | 82.74 |
| Asterids | Apiales | *Petroselinum crispum* | Parsley | Flat leaves | Botanical interests, Inc | 182 | 6.04 | 245.70 | 57.43 | 64.86 |
| Rosids | Cucurbitales | *Cucumis sativus* | Cucumber | National Pickling | Green garden products | 293 | 556.34 | 1052.82 | 119.96 | 138.14 |
| Rosids | Fabales | *Phaseolus vulgaris* | Common bean | French filet | Botanical interests, Inc | 386 | 3.13 | 550.86 | 60.26 | 64.23 |
| Rosids | Fabales | *Vigna unguicula* | Cowpea | IT97K-499-35 | USDA GRIN | | 285.21 | 362.68 | 57.97 | 72.93 |
| Rosids | Brassicales | *Brassica oleacera* | Kale | Kale Italian Nero Toscana | Botanical interests, Inc | 244 | 187.79 | 572.77 | 41.53 | 61.15 |
| Rosids | Brassicales | *Raphanus sativus* | Radish | Radish Cherry belle | Botanical interests, Inc | 427 | 380.28 | 666.67 | 26.64 | 32.17 |
| Rosids | Brassicales | *Arabidopsis thaliana* | Arabidopsis | Col-0 | | | 245.70 | 308.29 | 36.46 | 54.01 |

**Table S1: Eudicot species studied to test their surface-dependent interaction with *B. cinerea.*** In addition to the information on the seeds' origin, the average leaf thickness, stomata density and lesion area on each leaf surface are provided.

| Species | Genotype | Background | Pathway | Effect | Reference |
|---|---|---|---|---|---|
| Arabidospsis thaliana | Col-0 | | WT | | |
| Arabidospsis thaliana | Ler-0 | | WT | | |
| Arabidospsis thaliana | anac055 | Col-0 | Jasmonic acid signaling | Pathogen sensitivity | Bu et al 2008 Cell research |
| Arabidospsis thaliana | Coi1-16 | Col-0 | Jasmonic acid signaling | Pathogen sensitivity | Ellis & Turner 2002 |
| Arabidospsis thaliana | Npr1 | Col-0 | Salicylic acid signaling | Pathogen sensitivity | Cao et al. 1997 |
| Arabidospsis thaliana | pad4 | Col-0 | Salicylic acid signaling | Pathogen sensitivity | Zhou et al 1999 |
| Arabidospsis thaliana | tga3-2 | Col-0 | Salicylic acid signaling | Pathogen sensitivity | Miao et al. 1995 |
| Arabidospsis thaliana | tgg1/tgg2 | Col-0 | Myrosinase | Toxicity bomb | Barth & Jander 2006 |
| Arabidospsis thaliana | MYB28 | Col-0 | Aliphatic glucosinolates | Aliphatic glucosinolates | Sonderby et al. 2007 |
| Arabidospsis thaliana | MYB29 | Col-0 | Aliphatic glucosinolates | Aliphatic glucosinolates | Sonderby et al. 2007 |
| Arabidospsis thaliana | myb29 | Col-0 | Aliphatic glucosinolates | Aliphatic glucosinolates | Sonderby et al. 2010 |
| Arabidospsis thaliana | myb28/29 | Col-0 | Aliphatic glucosinolates | Aliphatic glucosinolates | Sonderby et al. 2010 |
| Arabidospsis thaliana | myb34/51 | Col-0 | Indolic glucosinolates | Indolic glcosinolates | Frerigmann et al. 2014 |
| Arabidospsis thaliana | cyp79b2/b3 | Col-0 | Indolic glucosinolates | Indolic glcosinolates | Mikkelson et al. 2003 |
| Arabidospsis thaliana | AT1G26420 (fox5) | Col-0 | Indolic glucosinolates | Hub gene in Arabidopsis-Botrytis coexp | Zhang et al. 2017 |
| Arabidospsis thaliana | pad3 | Col-0 | Camalexin | inducible phytoalexin | Zhou et al. 1998 |
| Arabidospsis thaliana | gGP1 | Col-0 | Indol-cyanogenic glycosides | inducible phytoalexin | Rajnak et al. 2015 |
| Arabidospsis thaliana | CYP71a12 | Col-0 | Indol-cyanogenic glycosides | inducible phytoalexin | Rajnak et al. 2015 |
| Arabidospsis thaliana | Cyp82C2 | Col-0 | Indol-cyanogenic glycosides | inducible phytoalexin | Rajnak et al. 2015 |

**Table S2: *A. thaliana* wild accessions and mutant lines used in this study.** Mutants are deficient in SA and JA defense signaling, aliphatic and indolic glucosinolates, putative cyanogenic glycosides, and camalexin biosynthesis.

| | | | Eudicot Experiment | | Arabidopsis Experiment |
|---|---|---|---|---|---|
| **Isolate** | **Origin** | **Host** | **Host specificity** | **Virulence** | **Camalexin sensitivity** |
| 2004 | California | Grape | | | |
| 01_01_01 | California | Grape | | | |
| 01_01_02 | California | Grape | | | |
| 01_01_03 | California | Grape | | | |
| 01_01_04 | California | Grape | | | |
| 01_01_06 | California | Grape | | | |
| 01_01_15 | California | Grape | | | |
| 01_02_01 | California | Grape | | | |
| 01_02_02 | California | Grape | | | |
| 01_02_03 | California | Grape | | | |
| 01_02_04 | California | Grape | | | |
| 01_02_05 | California | Grape | | | |
| 01_02_06 | California | Grape | | | |
| 01_02_15 | California | Grape | | | |
| 01_02_16 | California | Grape | | | |
| 01_02_17 | California | Grape | | | |
| 01_02_18 | California | Grape | | | |
| 01_02_20 | California | Grape | | | |
| 01_03_02 | California | Grape | | | |
| 01_03_04 | California | Grape | | | Medium |
| 01_03_12 | California | Grape | | | |
| 01_03_16 | California | Grape | | | |
| 01_03_18 | California | Grape | | | |
| 01_03_19 | California | Grape | | | |
| 01_03_20 | California | Grape | | | |
| 01_03_22 | California | Grape | | | |
| 01_03_23 | California | Grape | | | |
| 01_04_01 | California | Grape | | | |
| 01_04_02 | California | Grape | | | |
| 01_04_03 | California | Grape | | | |
| 01_04_04 | California | Grape | | | |
| 01_04_05 | California | Grape | | | |
| 01_04_12 | California | Grape | | | |
| 01_04_15 | California | Grape | | | |
| 01_04_17 | California | Grape | | | |
| 01_04_19 | California | Grape | | | |
| 01_04_20 | California | Grape | | | |

| | | | | | |
|---|---|---|---|---|---|
| 01_04_21 | California | Grape | | | |
| 01_04_25 | California | Grape | | | |
| 01_05_04 | California | Grape | | | |
| 01_05_11 | California | Grape | | | |
| 01_05_14 | California | Grape | | | |
| 01_05_16 | California | Grape | | | |
| 01_05_22 | California | Grape | | | |
| 01_05_24 | California | Grape | | | |
| 02_04_03 | California | Grape | | | |
| 02_04_04 | California | Grape | | | |
| 02_04_08 | California | Grape | | | |
| 02_04_09 | California | Grape | | | |
| 02_04_11 | California | Grape | | | |
| 02_04_12 | California | Grape | Medium | High | |
| 02_04_14 | California | Grape | | | |
| 02_04_17 | California | Grape | | | Medium |
| 02_04_18 | California | Grape | | | |
| 02_04_20 | California | Grape | | | |
| 02_04_21 | California | Grape | | | |
| 94_1 | California | Lemon | | | |
| 94_4 | California | Orange | | | |
| Acacia | South Africa | Acacia | | | |
| Apple404 | California | Apple | | | |
| Apple517 | California | Apple | Medium | Medium | High |
| Ausubel | Massachusetts | Brassica | | | |
| B05_10 | Netherlands | Grape | Low | Low | |
| BMM | Switzerland | Geranium | | | |
| BPA1 | California | Grape | | | |
| DavisNavel | California | Orange | High | Low | |
| EsparatoFresa | California | Strawberry | | | |
| Fd1 | California | Raspberry | | | |
| Fd2 | California | Raspberry | | | |
| Fresa525 | California | Strawberry | | | |
| FresaSD | California | Strawberry | | | |
| Gallo1 | California | Grape | | | |
| Gallo2 | California | Grape | | | |
| Geranium | California | Geranium | | | |
| Grape | South Africa | Grape | | | |
| KatieTomato | California | Tomato | Medium | High | Low |
| KernA2 | California | Grape | | | |
| KernB1 | California | Grape | Medium | Medium | Medium |

| | | | | | |
|---|---|---|---|---|---|
| KernB2 | California | Grape | | | |
| KGB1 | California | Tomato | | | |
| KGB2 | California | Tomato | | | |
| MEAP6G | Switzerland | Grape | | | |
| Mex03 | Mexico | Blackberry | | | |
| Molly | California | Grape | | | |
| Navel | California | Orange | | | |
| NobleRot | California | Grape | | | |
| Peachy | California | Peach | | | |
| Pepper | United Kingdom | Pepper | High | Low | Low |
| PepperSub | United Kingdom | Pepper | | | |
| PhiloMenlo | California | Grape | | | |
| Rasp | California | Raspberry | | | High |
| Rose | California | Rose | High | Low | Low |
| Supersteak | California | Tomato | | | |
| Triple3 | California | Tomato | Medium | Medium | Medium |
| Triple7 | California | Tomato | | | |
| UKRazz | United Kingdom | Raspberry | Medium | High | Very low |

**Table S3: Botrytis strains used in this study.** For each of the 96 strains the geographical origin and the host on which it was isolated are provided. For the 10 strains selected for the Eudicot and Arabidopsis experiments, the host specificity, virulence and camalexin sensitivity estimates are provided.

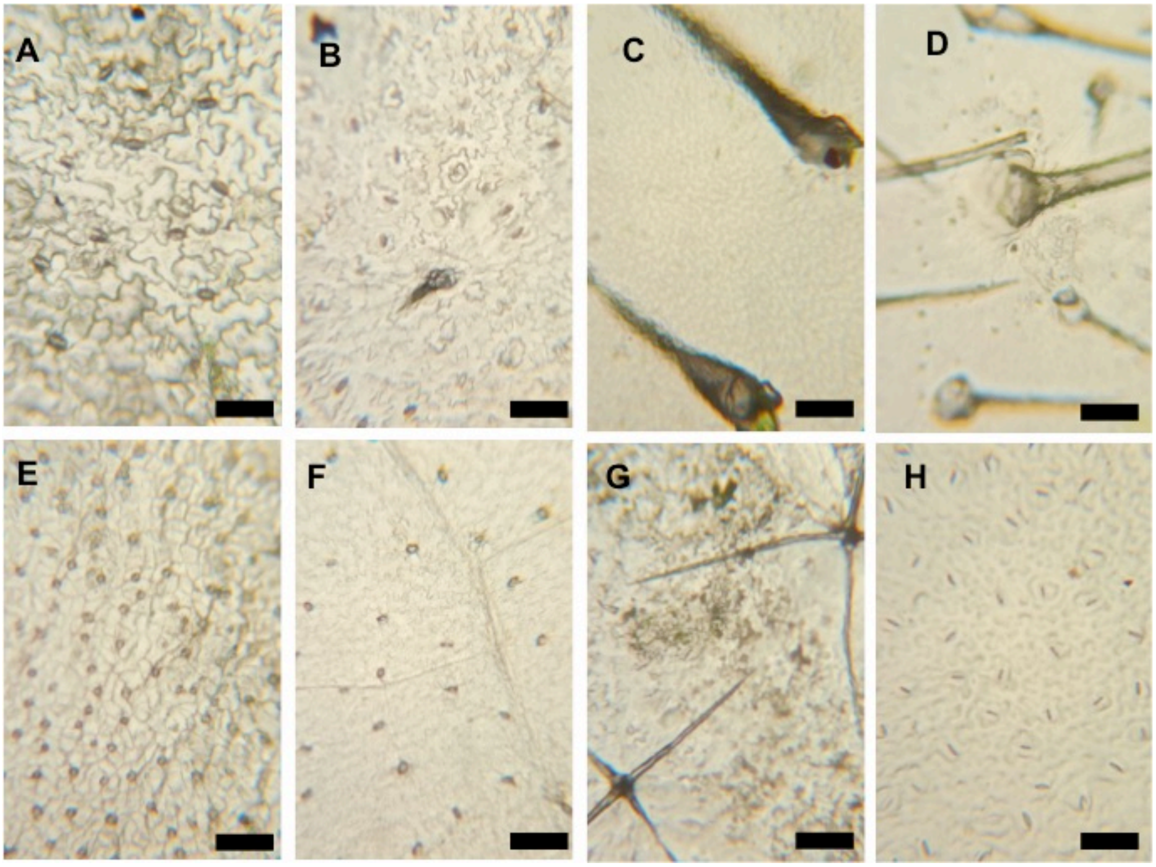


**Figure S1: Examples of the diversity of leaf surfaces among the eudicot species.** Stomata size varies greatly between species while the micro-topography is influenced by leaf shape, vein and trichome density. A) Adaxial surface of *Beta vulgaris* (chard) with pavement cells and stomata. B) Adaxial surface of *Ocimum basilicum* (basil) with pavement cells and stomata. C) Adaxial surface of *Solanum lycopersicum* (tomato) with linear trichome. D) Adaxial surface of *Cucumis sativus* (cucumber) with high density of linear trichomes. The microtopography of cucumber is highly uneven, with raised area around trichomes and valleys in between. E) Abaxial surface of *Spinacia oleracea* (spinach) with round stomata. F) Abaxial surface of *Lactuca sativa* (lettuce) with veins and stomata. G) Abaxial surface of *Solanum melongena* (eggplant) with high-branched trichomes and uneven microtopography. H) Abaxial surface of *Vigna unguiculata* (cowpea) with large stomata. The scale represents 100um. Pictures by Celine Caseys.

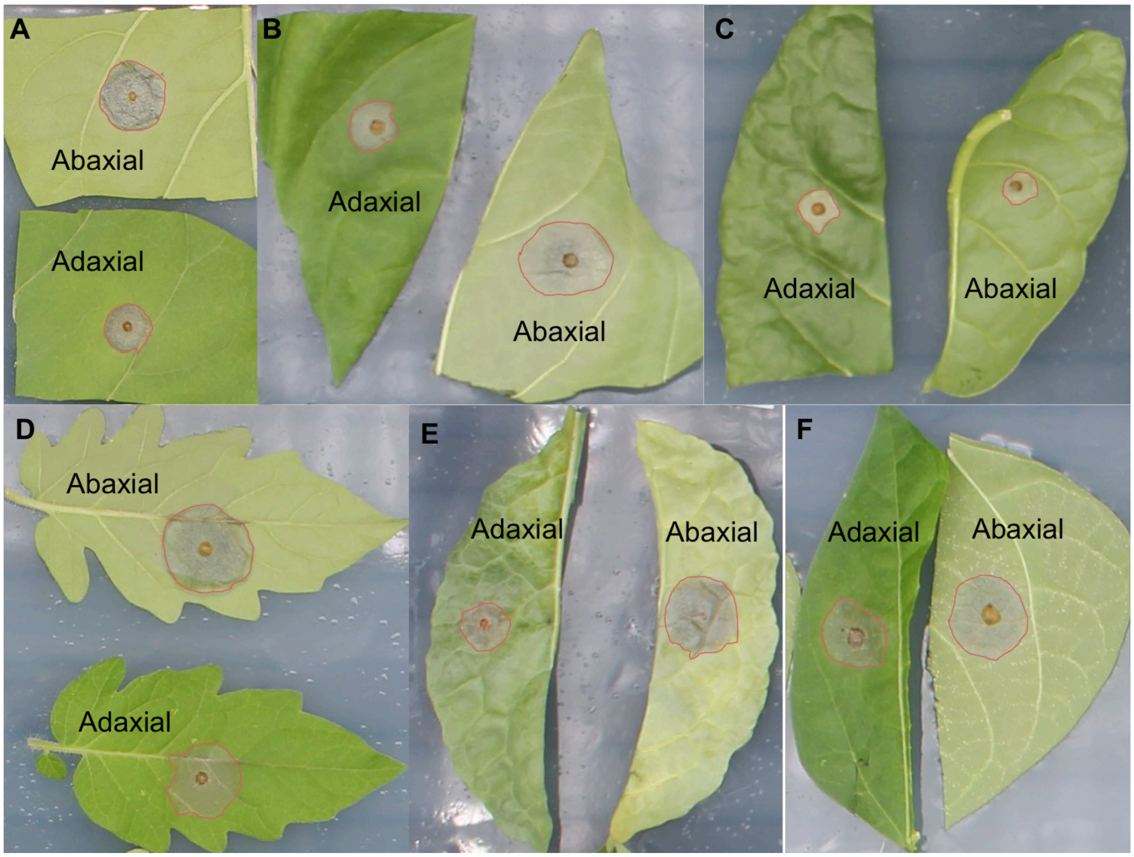


**Figure S2: Representative raw images of leaf abaxial and adaxial surfaces from detached leaf assay experiments**. On each surface, a red line highlights the lesion. A) *S.melongena* B) *O.basilicum* C) *B.vulgaris* D) *S.lycopersicum* E) *S.oleracea* F) *P. vulgaris.* Pictures by Celine Caseys.

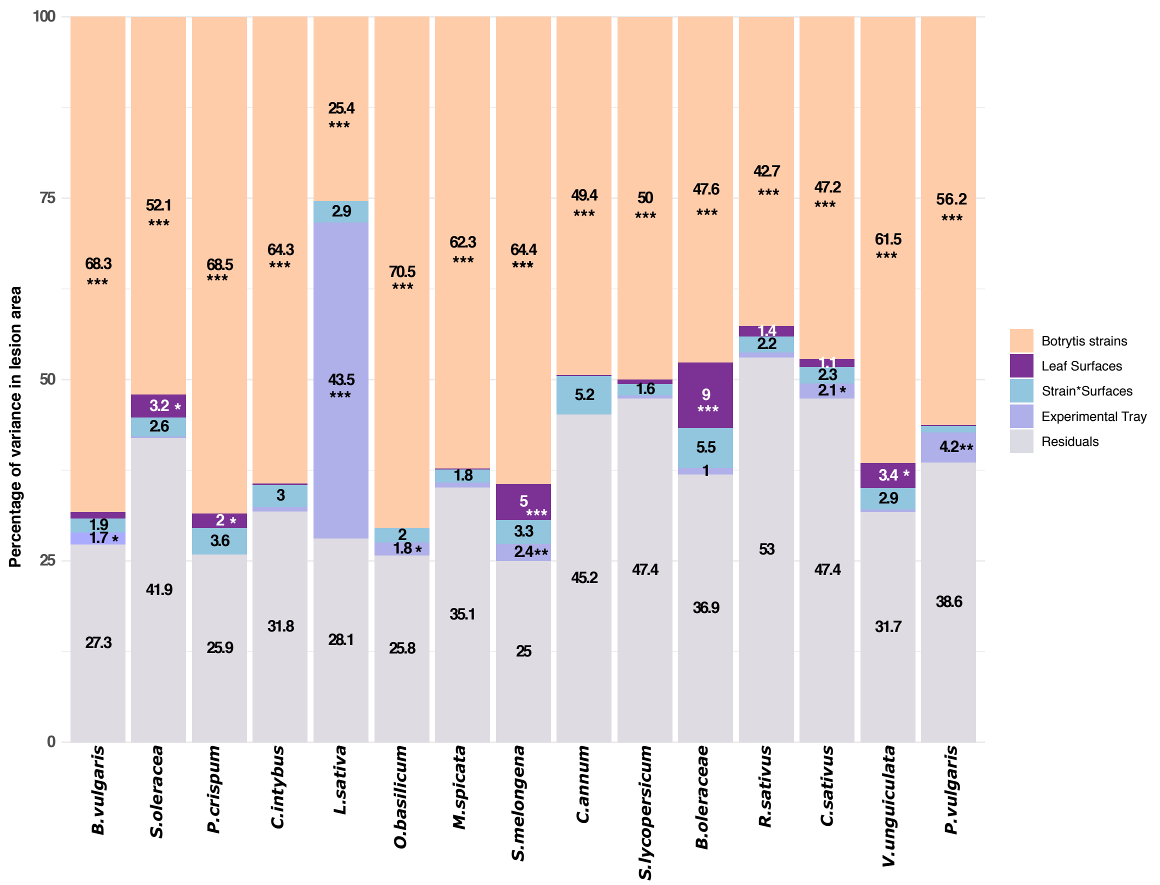


Figure S3: **Linear modeling of the lesion area on the Eudicot species.** The y-axis represents the percentage of variance explained by the Botrytis strain, the ab/ad-axial leaf surfaces, the interaction of the strain with the leaf surface and the experimental tray. Values for percentage of variance larger than 1% are provided. Significance: *** p<0.001; ** p<0.01; * p<0.05

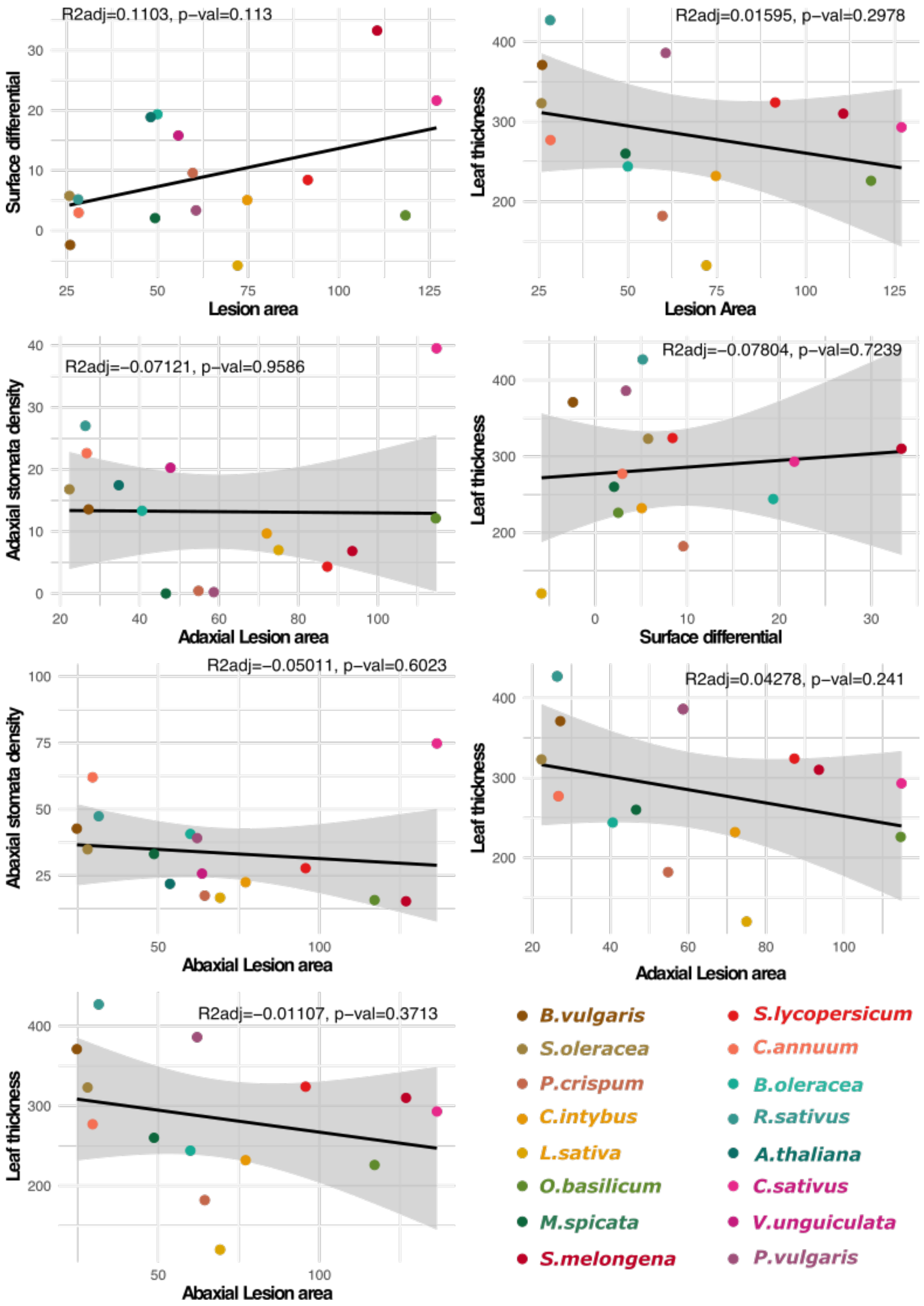


**Figure S4: Correlation between lesion area, stomatal density and leaf thickness.**

The back line represents the linear regression and the grey area the confidence interval. The adjusted $R^2$ and p-value of the linear regression are provided.

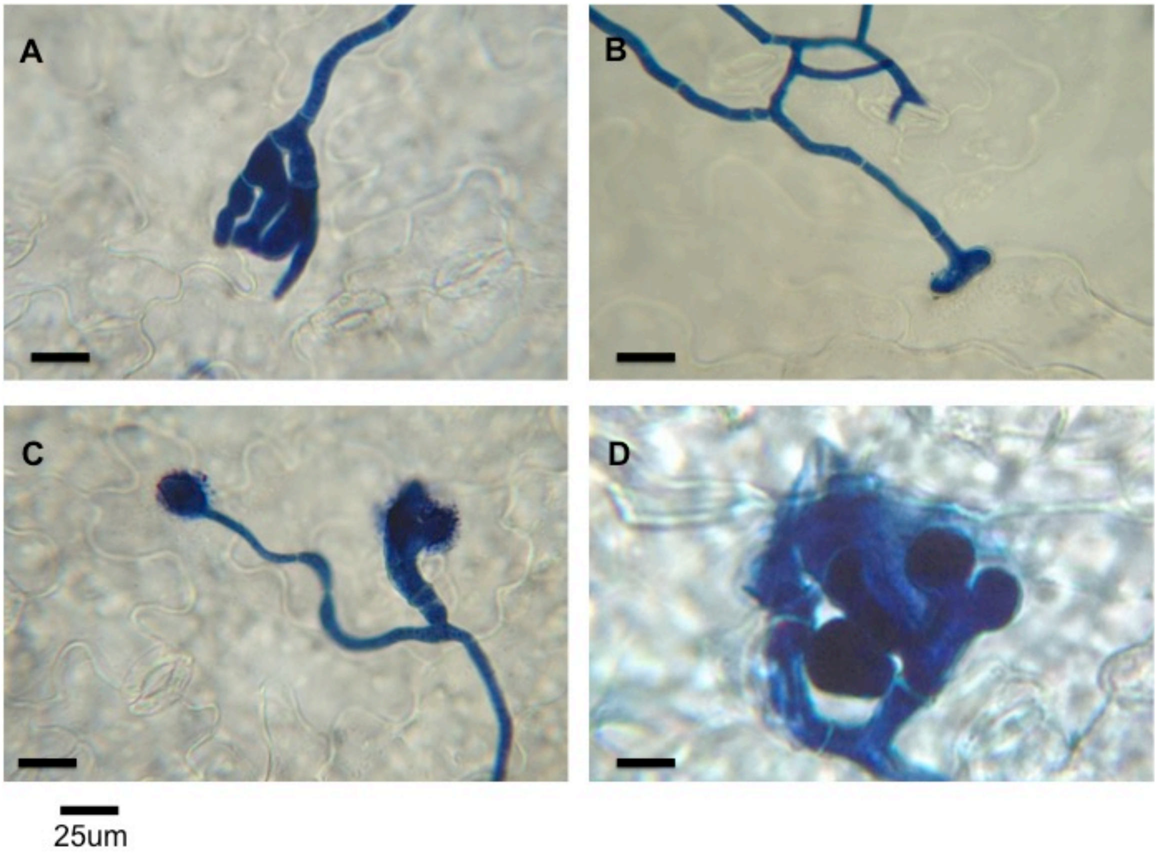


**Figure S5: Microscopy of *B. cinerea* stained with trypan blue on Col-0 leaf surfaces.** A) Branching hyphae on abaxial surface at 36 HPI. B) Hyphae on adaxial surface at 42 HPI. C) Infection cushions with extracellular vesicles on abaxial surface at 36 HPI. D) Infection cushion penetrating the adaxial surface at 48 HPI. The scale represents 25um. Pictured by Aleshia Hopper.

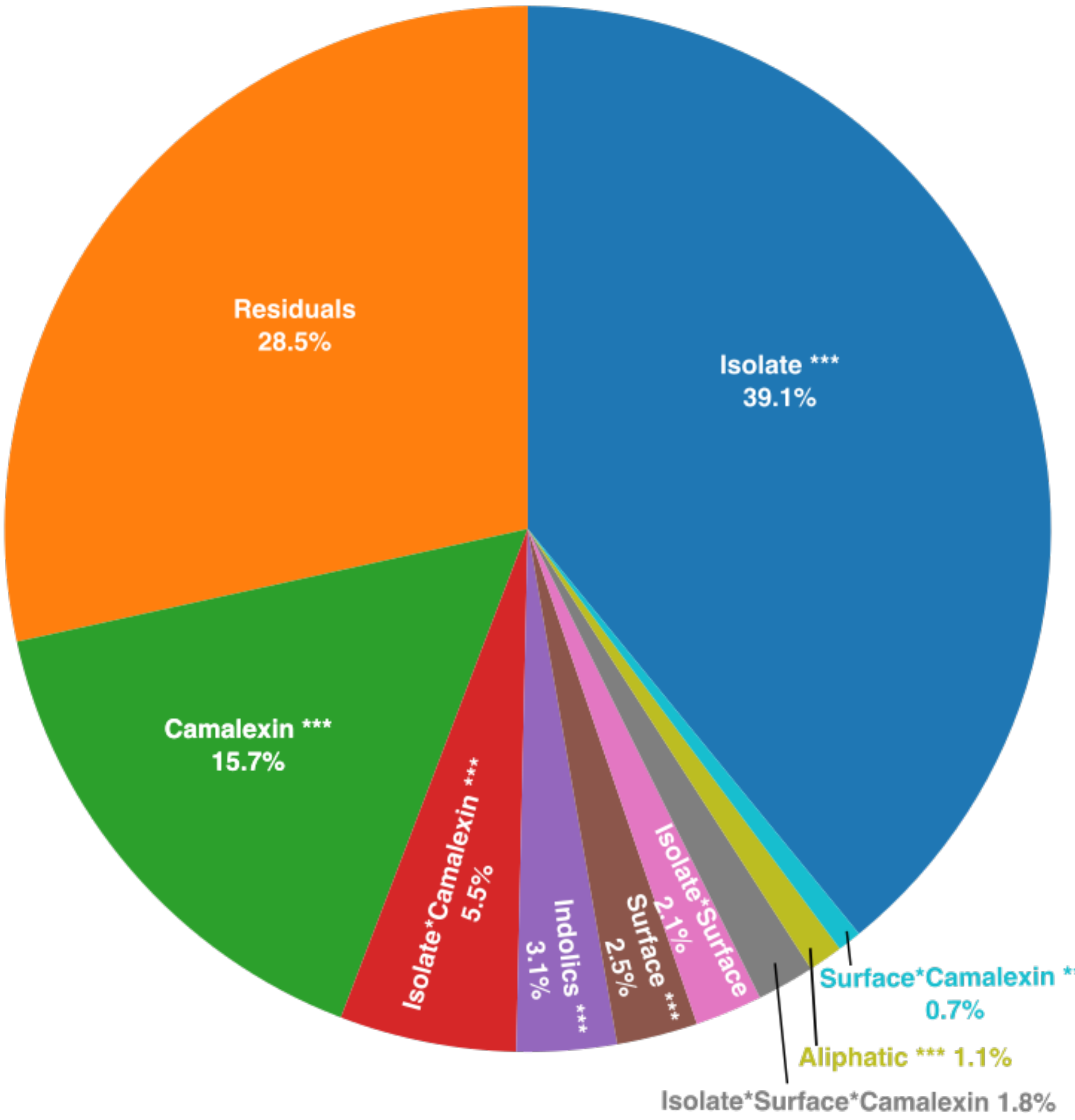


**Figure S6: Pie chart representing the percentage of variance in lesion area explained by the Botrytis isolates, camalexin, indolic glucosinolates, aliphatic glucosinolates and the leaf surface alone or in interaction estimated by linear modeling.** Significance: *** p<0.001; ** p<0.01; * p<0.05.

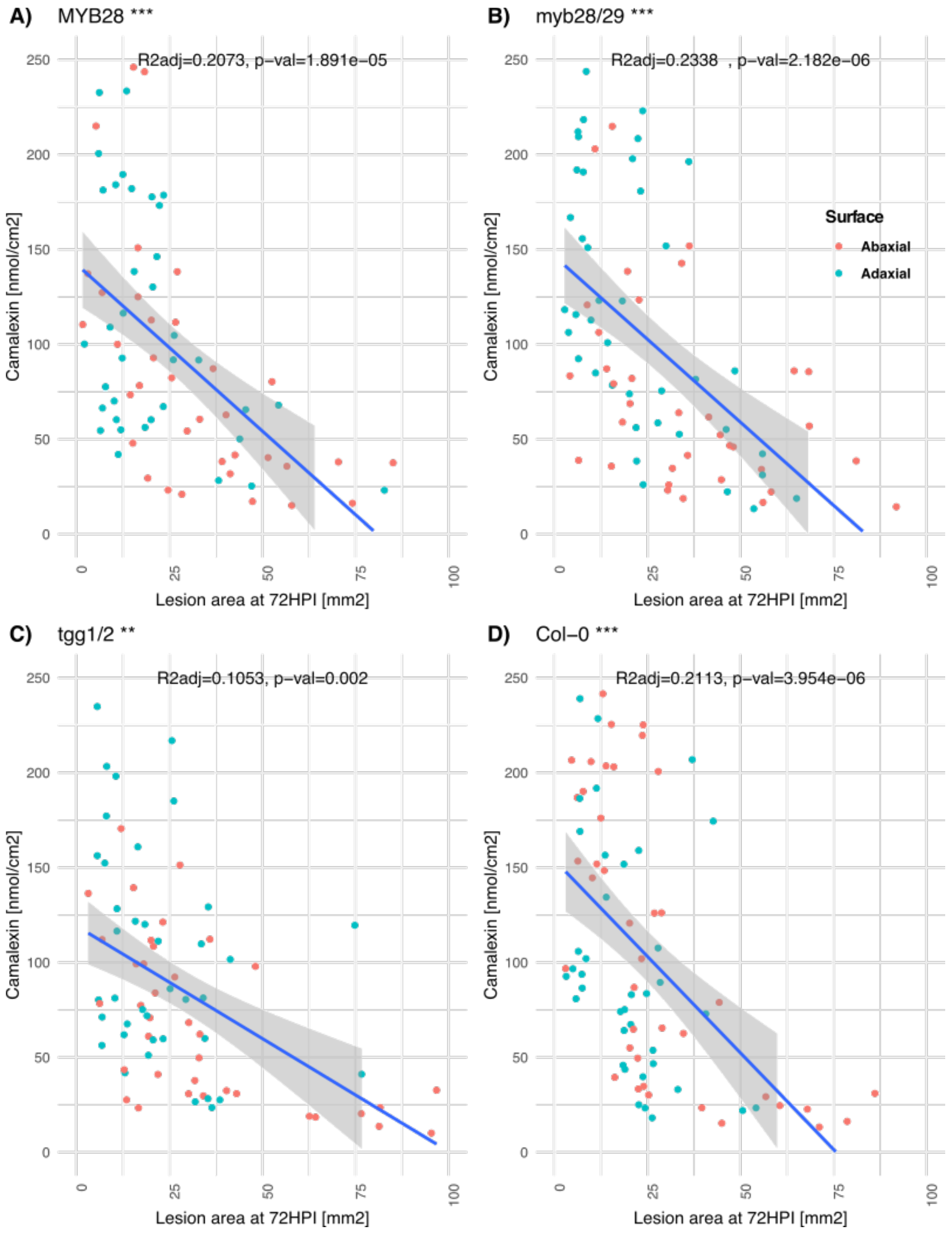


**Figure S7: Correlation between Botrytis growth on the leaf (lesion area) and concentration in camalexin produced by the plant.** Green dots represent Botrytis infections on the adaxial surface while red dots represent infections on the abaxial surface. The blue line represents the linear regression and the grey area the confidence interval. The adjusted $R^2$ and p-value of the linear regression are provided. Significance: *** $p<0.001$; ** $p<0.01$; * $p<0.05$

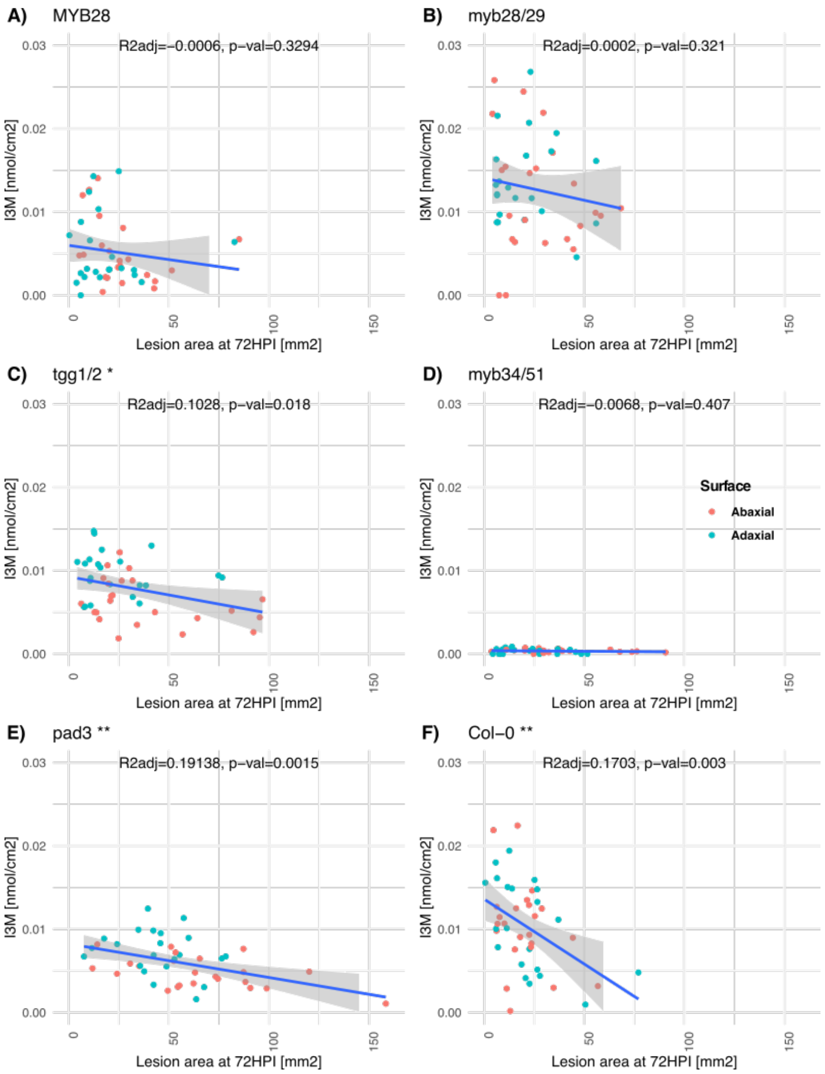


**Figure S8: Correlation between Botrytis growth on the leaf (lesion area) and concentration in indol-3-yl-methylglucosinolate (I3M) produced by the plant.** Green dots represent Botrytis infections on the adaxial surface while red dots represent infections on the abaxial surface. The blue line represents the linear regression and the grey area the confidence interval. The adjusted $R^2$ and p-value of the linear regression are provided. Significance: *** p<0.001; ** p<0.01; * p<0.05

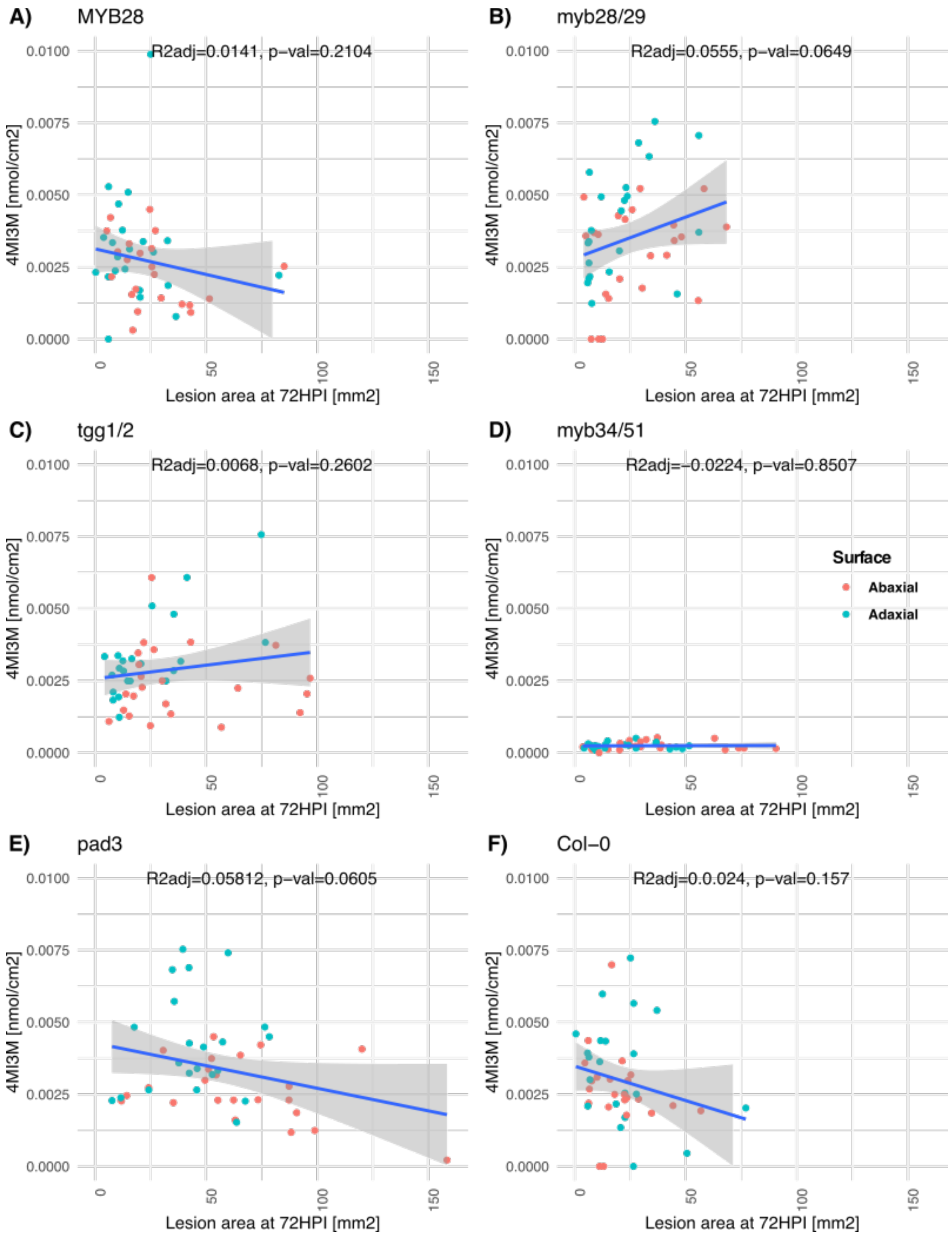


**Figure S9: Correlation between Botrytis growth on leaf (lesion area) and concentration in the 4-methoxy-indole-3-ylmethyl-GSL (4MI3M) produced by the plant.** Green dots represent Botrytis infections on the adaxial surface while red dots represent infections on the abaxial surface. The blue line represents the linear regression and the grey area the confidence interval. The adjusted $R^2$ and p-value of the linear regression are provided.

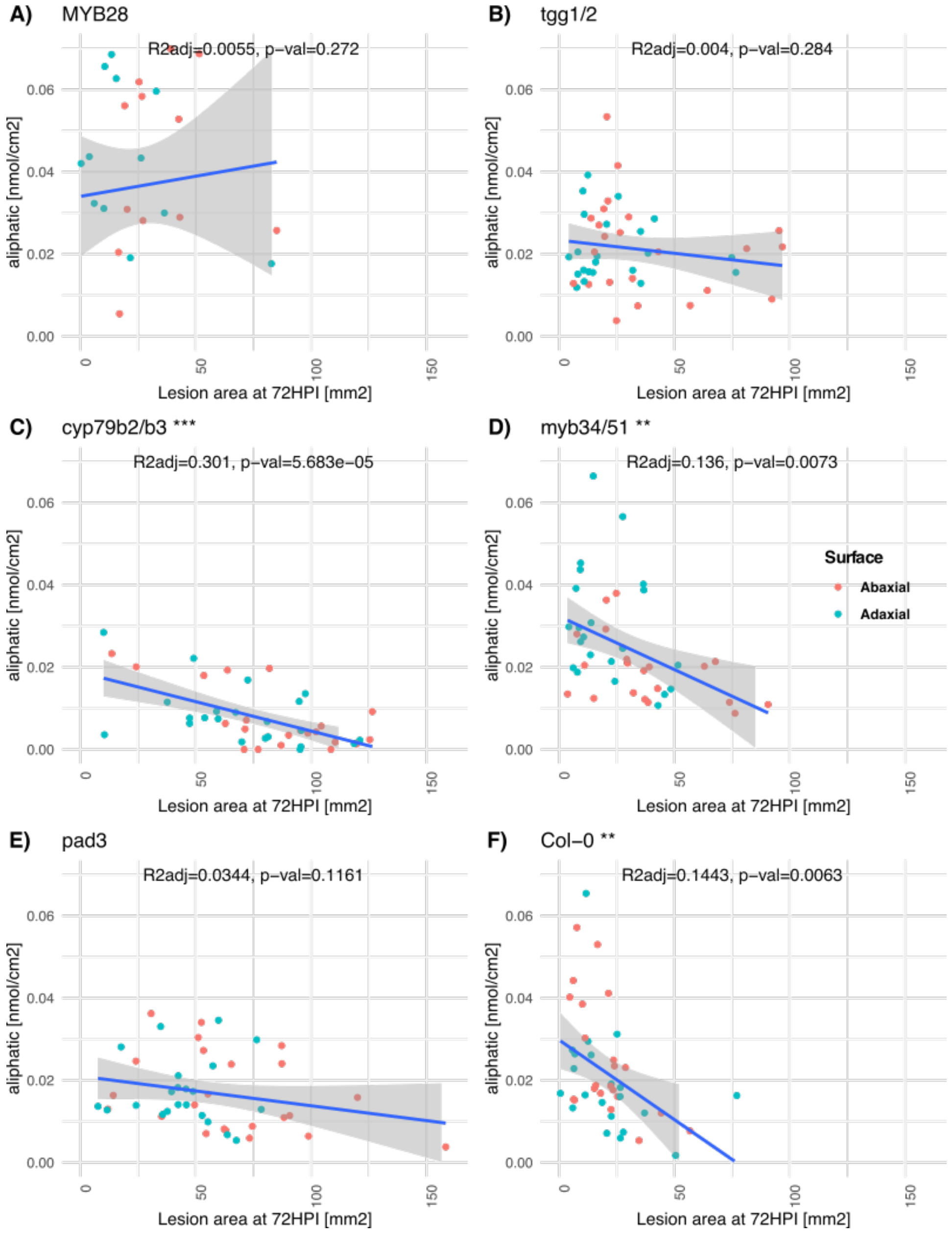


**Figure S10: Correlation between Botrytis growth on leaf (lesion area) and concentration in the aliphatic glucosinolates produced by the plant.** Green dots represent Botrytis infections on the adaxial surface while red dots represent infections on the abaxial surface. The blue line represents the linear regression and the grey area the confidence interval. The adjusted $R^2$ and p-value of the linear regression are provided. Significance: *** p<0.001; ** p<0.01; * p<0.05
